# Ultrasound-mediated delivery of novel tau-specific monoclonal antibody enhances brain uptake but not therapeutic efficacy

**DOI:** 10.1101/2021.10.18.464732

**Authors:** Rinie Bajracharya, Esteban Cruz, Jürgen Götz, Rebecca M. Nisbet

## Abstract

Tau-specific immunotherapy is an attractive therapeutic strategy for the treatment of Alzheimer’s disease and other tauopathies. However, targeting tau effectively remains a considerable challenge due to the restrictive nature of the blood-brain barrier (BBB), which excludes 99.9% of peripherally administered antibodies. We have previously shown that the delivery of tau-specific monoclonal antibody (mAb) with low-intensity scanning ultrasound in combination with intravenously injected microbubbles (SUS^+MB^) increases the passage of IgG antibodies into the brain. SUS^+MB^ transiently opens tight junctions to allow paracellular transport, but also facilitates transcellular transport, particularly for larger cargoes. However, therapeutic efficacy after enhanced brain delivery has not been explored. To assess whether ultrasound-mediated delivery of tau-specific mAbs leads to an enhanced therapeutic response, K369I tau transgenic K3 mice were passively immunised once weekly for 12 weeks with a novel mAb, RNF5, in combination with SUS^+MB^. While none of the treatment arms improved behaviour or motor functions in these mice, we found that both RNF5 and SUS^+MB^ treatments on their own reduced tau pathology, but, surprisingly, the combination of both (RNF5+SUS^+MB^) did not achieve an additive reduction in tau pathology. This was despite observing increased antibody penetration in the brain. Interestingly, a significant fraction of the antibody in the combination treatment was visualized in brain endothelial cells, suggesting that paracellular transport may not be the preferred uptake mechanism for RNF5. Taken altogether, more research is warranted to develop SUS^+MB^ as a delivery modality for anti-tau antibodies.

## Introduction

Tauopathies, such as Alzheimer’s disease (AD), are a group of neurodegenerative diseases in which abnormally phosphorylated tau aggregates into intracellular tau inclusions characteristic of any given tauopathy. In pre-clinical trials, immunotherapy has emerged as a valuable technique to reduce tau pathogenesis. However, in human clinical trials, it turned out to be challenging to achieve the primary endpoint outcomes. While there may be several reasons for this, we argue that achieving sufficient antibody concentration in the brain is a large determinant of therapeutic efficacy. Here, the blood-brain barrier (BBB) is a significant impediment to the passage of therapeutics into the central nervous system (CNS), with only 2% of peripherally administered small molecule drugs entering the brain [1]. For large molecule drugs, such as IgG antibodies, this diminishes to 0.1% and thus highlights the demand for novel approaches to increase the antibody concentration in the brain to facilitate increased target engagement [2, 3]. An emerging technology to enhance the passage of drugs across the BBB is the use of low intensity focused ultrasound to transiently open the BBB [4, 5]. In previous work, we have optimised a technique whereby repeated low-intensity ultrasound is applied in a scanning mode together with intravenous injection of microbubbles (scanning ultrasound with microbubbles, SUS^+MB^), allowing for safe and transient BBB opening across the whole brain [6-8]. Whilst pre-clinical studies performed by our lab and others have demonstrated that molecules of various sizes, as well as antibodies, are able to be delivered to the brain using ultrasound [9-13], therapeutic efficacy after enhanced brain delivery of a tau-specific mAb has not yet been assessed. Therefore, we aimed to investigate the therapeutic efficacy of our novel anti-tau mAb, RNF5, and whether its ability to reduce pathogenic tau could be enhanced by combining its delivery with SUS^+MB^ in K396I tau transgenic mice, a model of tauopathy that develops motor deficits from an early age [14]. To examine this, we performed a preclinical study with multiple arms in which mice were treated with either RNF5 alone, SUS^+MB^ alone, or the combination of both (RNF5+SUS^+MB^) once weekly for 12 weeks, assessing behavioural and motor functions together with biochemical and histological read-outs as a proxy for therapeutic efficacy.

## Materials and Methods

### Antibodies

Primary antibodies used for western blotting (WB), immunohistochemistry (IHC) and immunofluorescence (IF) in this study were as follows: RNF5 (In-house; WB: 1:10,000), RN235 (In-house; IF: 1:5,000), Tau-5 (Sigma-Aldrich; WB: 1:1,000), AT180 (pT231) and AT8 (p202/205) (Thermo Fisher) (WB: 1:1,000) (IHC: 1:500), IBA1 (Wako) (IHC and IF: 1:400), CD68 (Bio-Rad) (IF: 1:500), MAP2 (Abcam) (IF: 1:500), Lectin (Sigma-Aldrich) (Biotinylated, 1:250), GFAP (Dako) (IF: 1:500). The secondary antibodies used in this study were as follows: Goat anti-mouse Alexa Fluor 488 (Thermo) (IF: 1:500), Goat anti-rabbit Alexa Fluor 555 (Thermo) (IF: 1:500), polyclonal rabbit anti-mouse IgG biotinylated and polyclonal goat anti-rabbit IgG biotinylated (Dako) (IHC: 1:500), donkey anti-mouse IRDye 800CW, donkey anti-rabbit IRDye 680RD (LI-COR; WB: 1:10,000).

### Antibody generation

RNF5 is a monoclonal tau-specific antibody specific for amino acids 35-44 of human 2N4R tau. Large scale low-endotoxin antibody purification was conducted by the antibody services at the Walter and Elisa Hall Institute Biotechnology Centre. Final preparations in 1X PBS were stored in aliquots at a concentration of 8 mg/ml at -80°C. The isotype of RNF5 was determined using the IsoStrip Mouse Monoclonal Antibody Isotyping Kit (Sigma-Aldrich) and found to be IgG2b with a kappa light chain.

### Surface plasmon resonance

Surface plasmon resonance measurements were performed and analysed with assistance from the National Biologics Facility (NBF) at the University of Queensland. RNF5 was tested for binding to human 2N4R tau (hTau) and mouse 2N4R tau (mTau) using the Biacore T200 instrument. Briefly, hTau and mTau were immobilised onto an amine-coupling activated (NHS/EDC) CM5 Series S Sensor Chip (GE Healthcare) with a contact time of 180 s at 5 µL/min flow rate and at 25°C. For immobilisation, hTau was diluted to 42 nM and mTau to 455 nM in acetate buffer at pH 5.5 and pH 4, respectively. Binding of RNF5 to tau was tested using a Multi-Cycle High Performance Kinetics assay. RNF5 injections were performed using HPS-EP+ buffer (GE Healthcare) as a running buffer at a flow rate of 30 µL/min for 180 s followed by a dissociation time of 1,200 s in 3-fold increasing concentration steps (0.5-44.3 nM). For a given concentration, injections were done in duplicate, and regeneration of the chip surface was achieved with glycine pH 1.7. The data was blank-subtracted and fitted to a Two-State reaction model, using BIAevaluation software (GE Healthcare).

### Animals

All animal experiments were conducted following the guidelines of the Australian Code of Practice for the Care and Use of Animals for Scientific Purposes and approved by the University of Queensland Animal Ethics Committee (QBI/554/17/NHMRC). For treatment studies, 5-week-old male K3 mice expressing human 1N4R tau with the K369I mutation under the control of mThy.1.2 promoter were utilized [14]. Age-matched wild-type (WT) male littermates were used as a control in behavioural experiments. For labelled antibody delivery experiments, 10 – 30 week-old K3 mice of mixed gender were used.

### Production of microbubbles

Microbubbles comprising a phospholipid shell and octafluoropropane gas core were generated in-house as previously described [13]. Microbubble preparations yielded a mixture of microbubbles that were under 10 µm in size, with a mean diameter of 1 µm and a concentration of 1 – 5 × 10^9^ microbubbles/ml, similar to what has been reported for commercial microbubble preparations such as Definity^®^.

### SUS equipment

An integrated focused ultrasound system was used (TIPS, Philips Research) [6]. This system consisted of an annular array transducer with a natural focus of 80 mm, a radius of curvature of 80 mm, a spherical shell of 80 mm with a central opening of 31 mm diameter, a 3D positioning system, and a programmable motorized system to move the ultrasound focus in the x and y planes to cover the entire brain area. A coupler mounted to the transducer was filled with degassed water and placed on the head of the mouse with ultrasound gel for coupling, to ensure unobstructed propagation of the ultrasound to the brain.

### SUS and immunisation treatment

5-week-old K3 male and female mice were randomly assigned to one of the following groups: RNF5 only (RNF5), SUS^+MB^ only (SUS^+MB^), or SUS^+MB^ and RNF5 combined (RNF5+SUS^+MB^), with the inclusion of K3 sham and WT littermate controls for behavioural studies. All animals receiving injections and/or SUS^+MB^ treatment were anaesthetised and prepared as previously described [8]. Eight mice were used per experimental group and mice were treated once per week for 12 weeks. The sham K3 cohort received anaesthesia, but no retro-orbital injection or SUS^+MB^. For the antibody only and antibody plus SUS^+MB^ groups, 15 mg/kg of RNF5 was injected on its own via retro-orbital injection or as a mixture of 15 mg/kg of RNF5 and microbubbles (1 µL per gram body weight), respectively. Animals receiving SUS^+MB^ were placed under the ultrasound transducer, immobilised by a head frame. The parameters for the ultrasound delivery were 1 MHz centre frequency, 0.5 MPa peak rarefactional pressure (reduced from 0.7 MPa used in previous studies due to the young age and low weight of K3 mice [13]), 10 Hz pulse repetition frequency, 10% duty cycle, and a 6s sonication time per spot. The focus of the transducer was 1.5 mm and 12 mm in the transverse and axial planes, respectively. The motorized position system moved the focus of the transducer array in a grid with 1.5 mm spacing between individual sites of sonication such that ultrasound was delivered in a zig-zag pattern from the back of the brain moving towards the front of the brain. Mice received a total of 48 spots of sonication in a 6×8 raster grid pattern for the first 9 weeks of treatment, then received a total of 72 spots of sonication in a 9×8 raster grid pattern in the last 3 weeks of treatment to include sonication of the cerebellum.

### Behavioural tests

The weight of all animals was monitored weekly in both experiments involving K3 mice. Mice were tested weekly on the Rotarod to track motor function before, during and after treatment. Briefly, mice were habituated in the behavioural room for at least 30 minutes prior to testing. Light levels were kept constant at 60% (∼70 lux) throughout the test. Animals were placed on the Rotarod using the linear acceleration mode (4 to 40 rpm) over 90 s, with a total test time of 180 s (90 s at 40 rpm). Latency to fall was recorded as the longest time a mouse spent on the rod out of 5 attempts. Grip strength of forelimbs and hindlimbs was tested once before and once after antibody treatment by allowing mice to grasp a metal T-bar, and the average peak force to release the metal bar out of 10 trials was recorded. The activity monitor was used as an additional test to measure general animal behaviour before and after antibody treatment. The activity monitor is an automated animal tracking behavioural test whereby an individual animal is placed in a square-shaped open field box in which animal movement, position and speed are monitored by infrared beam breaks that project across the open field along the X, Y and Z axes. Mice were placed in the activity monitor box and allowed to explore for 30 minutes. The software was set up according to the General Open Field test from the Activity Monitor version 7 manual (MED Associates, Inc.). All data was transmitted to a PC and analysed using Activity Monitor 7 software, SOF-812 (MED Associates, Inc.). Untreated wild-type littermates were used as controls. Given that K3 mice exhibit a pronounced motor deficit, Jump time and Ambulatory time were included as read-outs.

### Tissue processing

24 hours after completion of the behavioural testing, treated mice and age-matched WT littermate controls were anaesthetised with an overdose of sodium pentabarbitone and transcardially perfused with PBS, their brains dissected, and the hemispheres separated. The right hemisphere was immersion-fixed in 4% PFA (Sigma) for 24 hours, and then embedded in paraffin using a benchtop tissue processor (Leica). Sagittal paraffin-embedded brain sections at 7 μm and 14 μm thickness were obtained using a microtome. The left hemisphere was snap-frozen in liquid nitrogen and stored at -80°C until further processing.

### Protein extraction

Snap-frozen brains from K3, WT and tau knockout mice were extracted using the sarkosyl extraction method with some modifications [15]. Briefly, snap-frozen hemispheres were homogenized in 3 volumes of RIPA-A buffer (50 mM Tris-HCl, pH 7.6, 150 mM NaCl, 1% Nonidet P-40, 5 mM EDTA, pH 8.0, 0.5% sodium deoxycholate, 50 mM sodium fluoride, 200 mM phenylmethyl sulfonyl fluoride and 100 mM sodium vanadate, supplemented with cOmplete™ mini protease inhibitor cocktail tablets (Roche)). Homogenates were centrifuged at 20,000 x g for 20 minutes at 4°C and the supernatant was transferred to a new tube and an equal volume of RIPA-B buffer (same as RIPA-A with the addition of 2% sarkosyl) was added to each. This was incubated for 1 hour at room temperature while rotating before centrifugation at 100,000 x g for 90 minutes at 4°C using the Optima™ MAX-XP Ultracentrifuge (Beckman Coulter). The supernatant was then collected (sarkosyl-soluble fraction). The remaining pellet was re-suspended in 250 μL RIPA-C buffer (same as RIPA-A with the addition of 1% SDS), sonicated and spun at 1000 x g for 1 minute to remove bubbles. Samples were then collected (sarkosyl-insoluble fraction). For both extraction methods, total protein concentration was estimated by performing a BCA assay (Thermo Fisher Scientific), and absorbance was measured at 562 nm using a POLARstar OPTIMA plate reader (BMG Labtech).

### Western blot analysis

5 μg of total protein was electrophoresed on a 10% Tris-glycine SDS-PAGE gel. Proteins were transferred to Immun-Blot Low Fluorescence PVDF membrane (Bio-Rad) using the Transblot Turbo Transfer System (Bio-Rad) and then stained with REVERT™ 700 Total Protein Stain according to the manufacturer’s instructions (LI-COR). Total protein was imaged in the 700 nm channel with the Odyssey FC Imaging System (LI-COR) then membranes were washed and blocked for 1 hour in Odyssey^®^ Blocking Buffer (LI-COR). Membranes were incubated in primary antibody solution overnight at 4°C by rocking. Membranes were washed with TBS-T and incubated with the secondary antibody for 1 hour at room temperature. Membranes were imaged in the 800 nm channel and fluorescence was quantified using the Image Studio™ Lite software (LI-COR).

### Labelled antibody delivery

RNF5 was covalently conjugated to Alexa Fluor 647 dye (Thermo Fisher Scientific) in PBS with 0.1M sodium bicarbonate, separated from free-dye and concentrated to volumes appropriate for injections, as described previously [9].

To compare antibody localisation of the RNF5 with and without SUS^+MB^, K3 mice of mixed gender aged 10 – 30 weeks old were treated with labelled RNF5 only (n=1 per time-point), SUS^+MB^ only (n=1 per time-point) or labelled RNF5 + SUS^+MB^ (n=3 per time-point) and perfused both 2 and 8 hours after treatment. Due to Covid-related unavailability of K3 mice at the time, only one mouse could be used per time-point in the RNF5 only and SUS^+MB^ only group. Mice were sonicated as described above. At their respective time-points, mice were transcardially perfused with PBS and their whole brains fixed in 4% PFA (Sigma) for immunohistochemical analysis.

### Antibody localisation assessment

To assess brain uptake of Alexa Fluor 647-conjugated RNF5, mouse brains were imaged using the Odyssey^®^ Fc Imaging System (700 nm laser, 2 minute scan time). Following this scan, whole brains were embedded in paraffin and sectioned coronally at 7 µm thickness using a microtome, followed by immunohistochemistry.

### Immunofluorescence and immunohistochemistry

Mounted brain tissue was first dehydrated in a series of xylene and ethanol washes. Antigen retrieval was conducted in a domestic microwave (850W) in citrate buffer pH 5.8 for 15 min and was cooled to room temperature. Sections were analysed by immunohistochemistry as described [16], using 2-3 sections per mouse. Briefly, sections were incubated with primary antibodies overnight. For immunofluorescence staining, sections were washed and incubated with respective secondary antibodies, then counterstained with DAPI nuclear stain (4’, 6-diamidino-2-phenylindole) (1:5,000). Alexa Fluor 647-conjugated RNF5 was detected *in situ* without additional amplification. For immunohistochemical staining, sections were stained using the nickel-diaminobenzidine (Nickel-DAB) method with no counterstain applied. All images were obtained on an automated slide scanner (Axio Imager Z2, Zeiss) using a Metafer Vslide Scanner program (Metasystems) at 20x magnification. Confocal Z-stack images were obtained on a Zeiss 510 confocal microscope at 63x magnification.

### Image analysis

Image quantification and analysis were performed blinded using ImageJ software. For all analyses, a control with no primary antibody was used for thresholding. More specifically, for immunofluorescence staining quantification, the ‘percentage (%) area positivity’ for RN235-, AT8- and AT180-tau was obtained on thresholded images of the cortex and hippocampus using the ‘area fraction’ method. For Nickel-DAB staining of Iba1, ‘Iba1-positive cell count’, ‘average cell size’ and ‘% immunoreactive area (area fraction method)’ were obtained by using the Analyse Particles plugin on thresholded images of an area of the primary somatosensory cortex. The regions of interests (ROIs) drawn in the Iba1 analyses were kept constant across images. Immunofluorescence staining quantification of ‘% area positivity’ for CD68 was obtained by using the ‘area fraction’ method. For analysis of ‘percentage (%) eosinophilic cells’ within the pyramidal layer of the hippocampus, images stained with hematoxylin and eosin were deconvoluted using the Colour Deconvolution 1.7 (H+E) plugin. Hematoxylin- and eosin-positive cell counts were obtained using the Analyse Particles plugin on thresholded images. ‘Percentage (%) eosinophilic cells’ was calculated by dividing total number of eosin-positive cells with total number of hematoxylin-positive cells. All measurements were noted and averaged over 2-3 sections per mouse.

To assess neuronal health of mice that were administered RNF5-647 in the presence of ultrasound, ‘% area positivity’ for MAP2 and Iba1 was quantified in areas that had RNF5 uptake versus areas that did not have RNF5 uptake using the ‘area fraction’ method on thresholded images. ROIs were manually drawn around areas that had RNF5 uptake and the same ROI was then used to quantify MAP2 and Iba1 in areas that did not have RNF5 uptake. For the per neuron correlative analysis between MAP2 and RNF5, the median fluorescence intensity of 63x confocal Z-stack images was analysed. Background levels of RNF5 fluorescence from a ‘no primary antibody control’ was measured and subtracted from the experimental data. MAP2 was used to draw ROIs around individual neurons. Neurons that were positive for RNF5, however, had indistinguishable MAP2 staining, therefore, for these neurons, RNF5-647 was used to draw ROIs around individual neurons.

### Statistical analysis

Statistical analyses were performed using GraphPad Prism 9.0 software. All data was analysed using one-way ANOVA followed by Tukey’s multiple comparisons test or t-test where appropriate, except for the Rotarod test where a two-way ANOVA was used, followed by Tukey’s multiple comparisons test. Correlations were assessed using the Spearman’s correlation test. All values are reported as mean ± standard error of the mean (SEM). Outliers were removed using the ROUT method (Q=1).

## Results

### RNF5 is a pan-tau monoclonal specific for amino acids 35-44

We generated a pan-tau antibody, RNF5, against a region within the N-terminus of tau spanning amino acids 35-44 (Figure 1A). To characterise the RNF5 IgG, we first established the specificity of RNF5 by western blotting with mouse brain lysates. This demonstrated that RNF5 binds total tau in the brain lysates of both wild-type and K3 tau transgenic mice that express 0N4R tau with the K369I mutation, thus indicating that RNF5 binds both mouse and human tau (Figure 1B). Western blot analysis of cortical brain homogenate from AD brains and healthy controls demonstrates that RNF5 can bind multiple forms of tau, further confirming that RNF5 is a pan-tau antibody (Figure 1C). Furthermore, in cortical brain tissue, RNF5 binds to the characteristic neurofibrillary tangles in AD brains (indicated by yellow arrow heads) which is absent from healthy controls (Figure 1D). Surface plasmon resonance (SPR) revealed that RNF5 has an affinity of 159 nM to 2N4R mouse tau and 25.3 nM to human 2N4R tau (Figure 1E and F). Thus, in vitro, RNF5 binds tau with 6-fold higher affinity than mouse tau (Figure 1G). Importantly, this data was fitted to a two-state reaction model as both the association and dissociation phases appear to have two components – a fast association followed by a slower association, and a fast dissociation followed by a slower dissociation.

**Figure 1:**
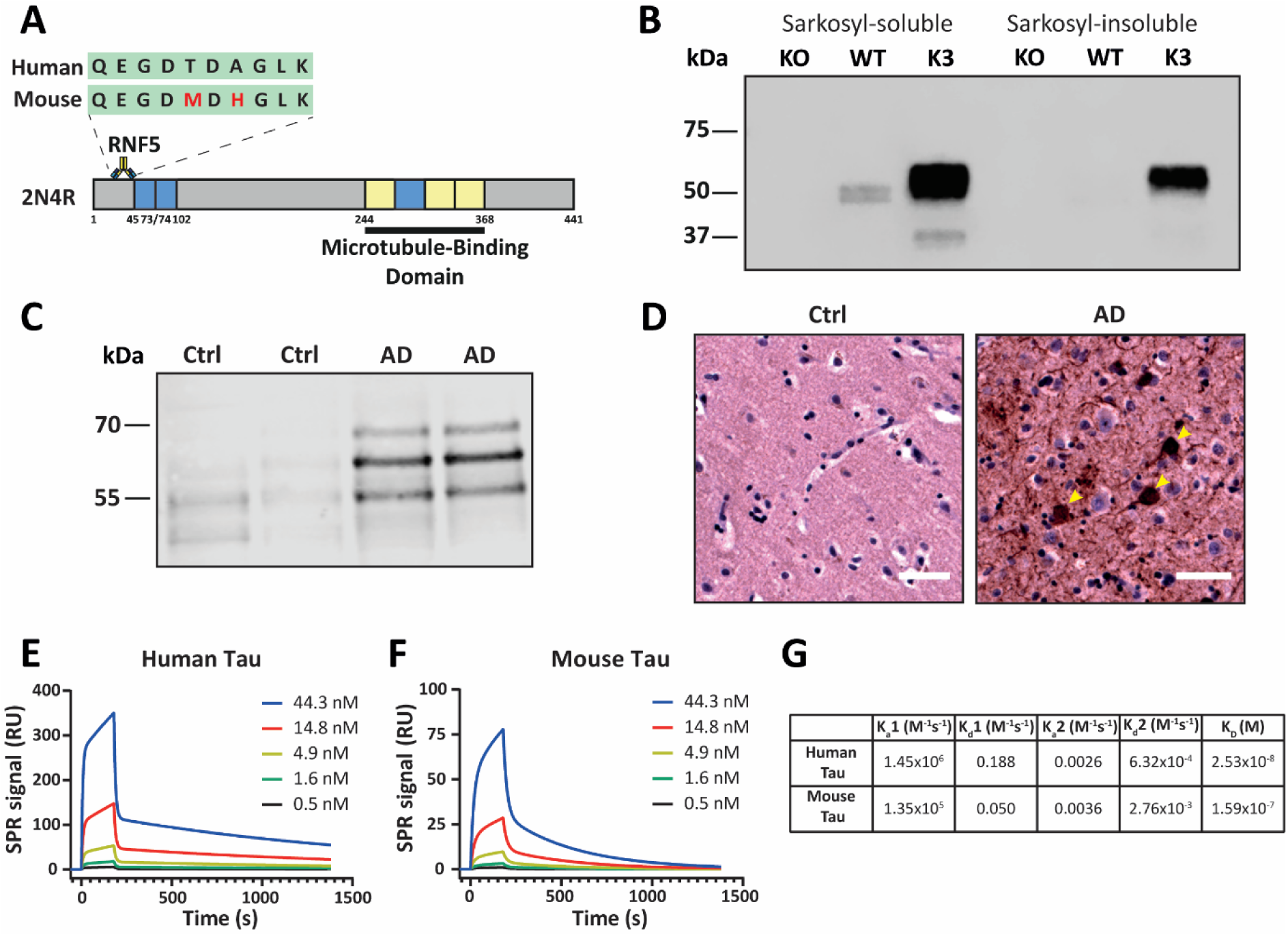
Characterisation of RNF5 by western blotting and SPR analysis. **(A)** Schematic representation of RNF5 bound to tau at amino acids (aa) 35-44. Target epitope (human and mouse amino acid sequences) of RNF5 is highlighted (green) with differing amino acids in red. **(B)** Western blot analysis of tau knockout (KO), wild-type (WT) and K369I tau transgenic mouse brain homogenate probed with RNF5 antibody. **(C)** Western blot analysis of healthy control (Ctrl) and Alzheimer’s disease (AD) cortical brain homogenate probed with RNF5 antibody. **(D)** DAB staining of cortical tissue from healthy control (ctrl) and AD brain probed with RNF5 antibody. Scale bar, 50 µm. **(E and F)** SPR sensograms of RNF5 antibody binding to immobilised human tau **(E)** and immobilised mouse tau **(F). (G)** Summary table showing association rate constant (K_a_), dissociation rate constant (K_d_), and binding constant (K_D_) of RNF5 against human and mouse tau. Kinetic parameters were calculated using a Two-State Reaction model.

### Neither RNF5, SUS^+MB^ nor the combination treatment improve the motor deficits of tau transgenic K369I mice

To investigate the therapeutic efficacy of RNF5 and whether its ability to reduce tau pathology could be enhanced by increasing its concentration in the brain, we combined the intravenous delivery of RNF5 in mice with SUS^+MB^ (Figure 2A). Mice treated with either RNF5 alone or SUS^+MB^ alone were used as controls, while litter-mate wild-type (WT) mice were used as additional control to the behavioural analysis. As reported previously, the weight of the K3 mice in this study was reduced compared to WT mice [14]. There was no significant difference, however, between the weight of treated mice compared to sham mice (Figure 2B). Weekly assessment of motor behaviour on the Rotarod revealed no significant improvements in latency to fall off the Rotarod for any of the treated groups compared to sham mice (Figure 2C). All treated mice, as well as the sham controls, improved at similar rates between 8 to 12 weeks of age. While the Rotarod can be used to test motor ability, coordination and limbic strength, we decided to also test the K3 mice in the activity monitor, to assess rearing ability (vertical time) and ambulation (ambulatory time), as well the grip strength test, to assess forelimb and hindlimb strength. K3 mice performed significantly worse in all tests compared to WT mice (Figure 2D to G). End-point behavioural testing after completion of the treatment revealed no significant improvements in rearing ability, ambulation or grip strength between any of treatment groups compared to sham (Figure 2D to G).

**Figure 2.**
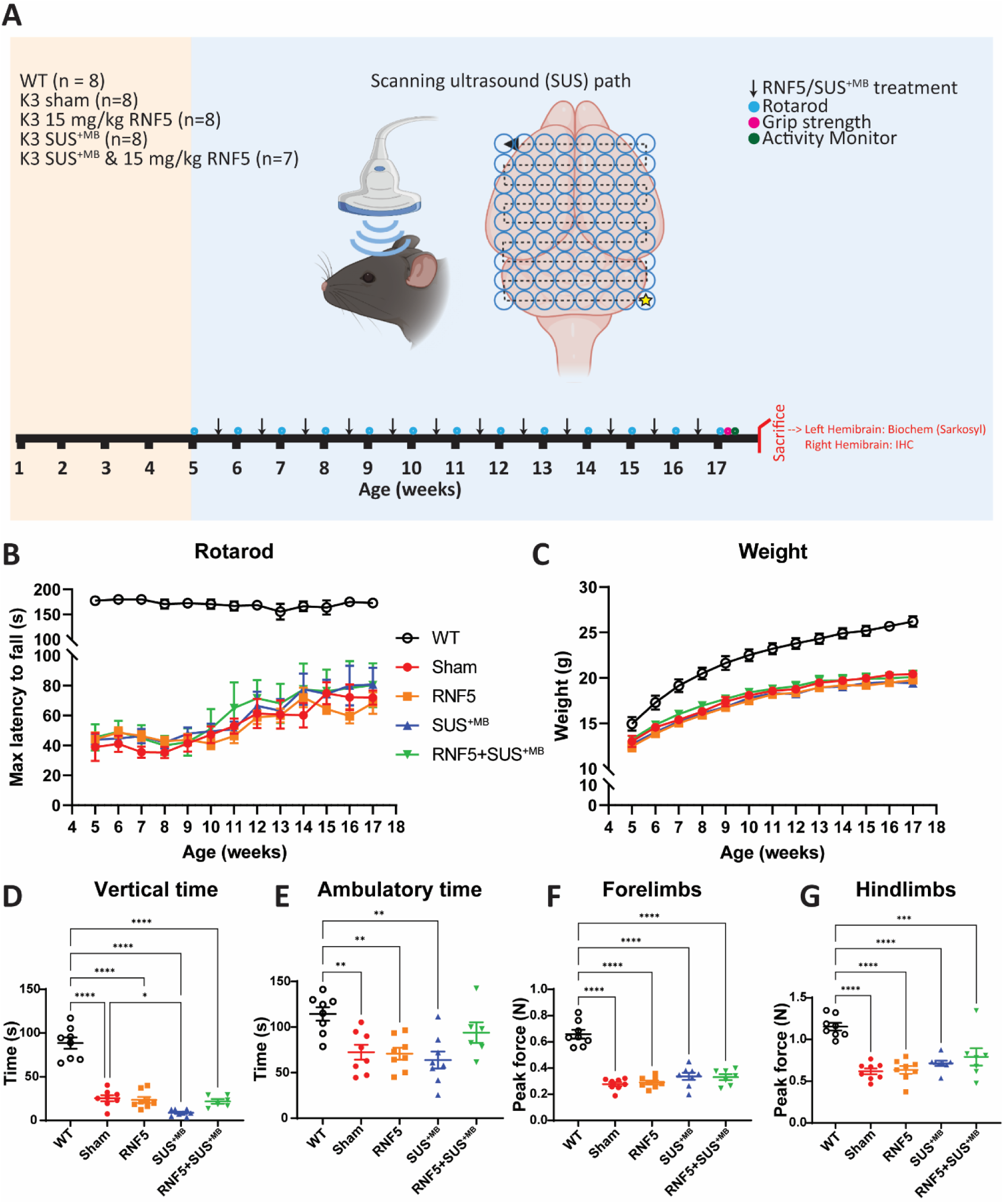
Motor ability and coordination is not improved following treatment with RNF5, SUS^+MB^ or RNF5+SUS^+MB^. **(A)** Schematic of treatment schedule with behaviour: 5-week-old male K3 tau transgenic mice were randomly assigned to one of four groups: Sham, RNF5, SUS^+MB^ and RNF5+SUS^+MB^. WT mice were included as behavioural controls. The scanning ultrasound (SUS) path indicates the 9×8 raster grid pattern of the ultrasound beam that was applied on mice receiving SUS^+MB^ with a yellow star representing the starting point (created with BioRender.com). **(B)** Mean latency to fall from the Rotarod for all four groups up to 17 weeks of age. There were no differences in latency to fall between groups **(C)** Mean weight of mice for all four groups up to 17 weeks of age (WT, n=8; Sham, n=8; RNF5, n=8; SUS^+MB^, n=8; RNF5+SUS^+MB^, n=7). **(D-G)** Behavioural testing performed after completion of treatment showed no differences between treated mice and Sham mice: **(D)** Vertical time (Activity Monitor) and **(E)** Ambulatory time (Activity Monitor), **(F)** Forelimb grip strength, **(G)** Hindlimb grip strength, All data represented as mean ± SEM (WT, n=8; Sham, n=8; RNF5, n=8; SUS^+MB^, n=8; RNF5+SUS^+MB^, n=7; ****p<0.0001, ***p<0.001, **p<0.01, *p<0.05, one-way ANOVA with Tukey’s multiple comparison’s test).

### All treatment arms reduce levels of phosphorylated tau in cortical and hippocampal tissue of K3 mice

We next sought to investigate whether the treatment had any effect on total and phosphorylated tau species in K3 mice. The strain develops robust tau pathology particularly in the cortex and hippocampus, and to a lesser extent in the cerebellum [14]. As pathology is most robust in the former two areas, the cerebellum was not analysed. Firstly, using immunofluorescent analyses, we assessed 3 different phospho-tau antibodies, RN235, AT8 and AT180, all of which stain tau inclusions to varying degrees [17] and bind to epitopes of tau that are elevated in AD [18, 19]. Labelling with RN235, which detects Pick body-like inclusions in K3 mice and is specific for phospho-S235, revealed significant reductions (by approximately 2-fold) in the number of RN235-positive inclusions in both the cortex and hippocampus of treated mice, in comparison to sham mice and no significant difference between each of the treated groups (Figure 3A and 4A). Labelling with AT8, which detects both Pick body-like inclusions and diffuse species of tau in K3 mice and is specific for phospho-S202/T205, revealed a significant reduction in the AT8-positive area in the cortex of the SUS^+MB^ only (by 19-fold) and RNF5 only (by 5-fold) treated groups but surprisingly, not in the RNF5+SUS^+MB^ treated group, compared to sham (Figure 3B). In the hippocampus, however, the AT8 positive area was significantly reduced only in the SUS^+MB^-treated mice (by 2.5-fold) compared to sham mice, as well as compared to RNF5- and combination-treated mice (Figure 4B). Finally, labelling with AT180, which predominantly detects pre-NFT-like inclusions and diffuse species of tau in K3 mice and is specific for phospho-T231, revealed significant reductions in the AT180-positive area in the cortex for all treatment groups (RNF5, SUS^+MB^ and RNF5+SUS^+MB^) compared to sham mice (by approximately 2.5-fold), with no significant difference between RNF5 and RNF5+SUS^+MB^ treated mice (Figure 3C). In the hippocampus, there was a trend towards a reduction in AT180-positive area across all treatment groups in comparison to sham mice (Figure 4C), however this did not reach significance. Taken altogether, these results indicate that while RNF5, SUS^+MB^ and the combination of RNF5+SUS^+MB^ are all successful at reducing tau pathology, the combination treatment provided no added benefit.

**Figure 3.**
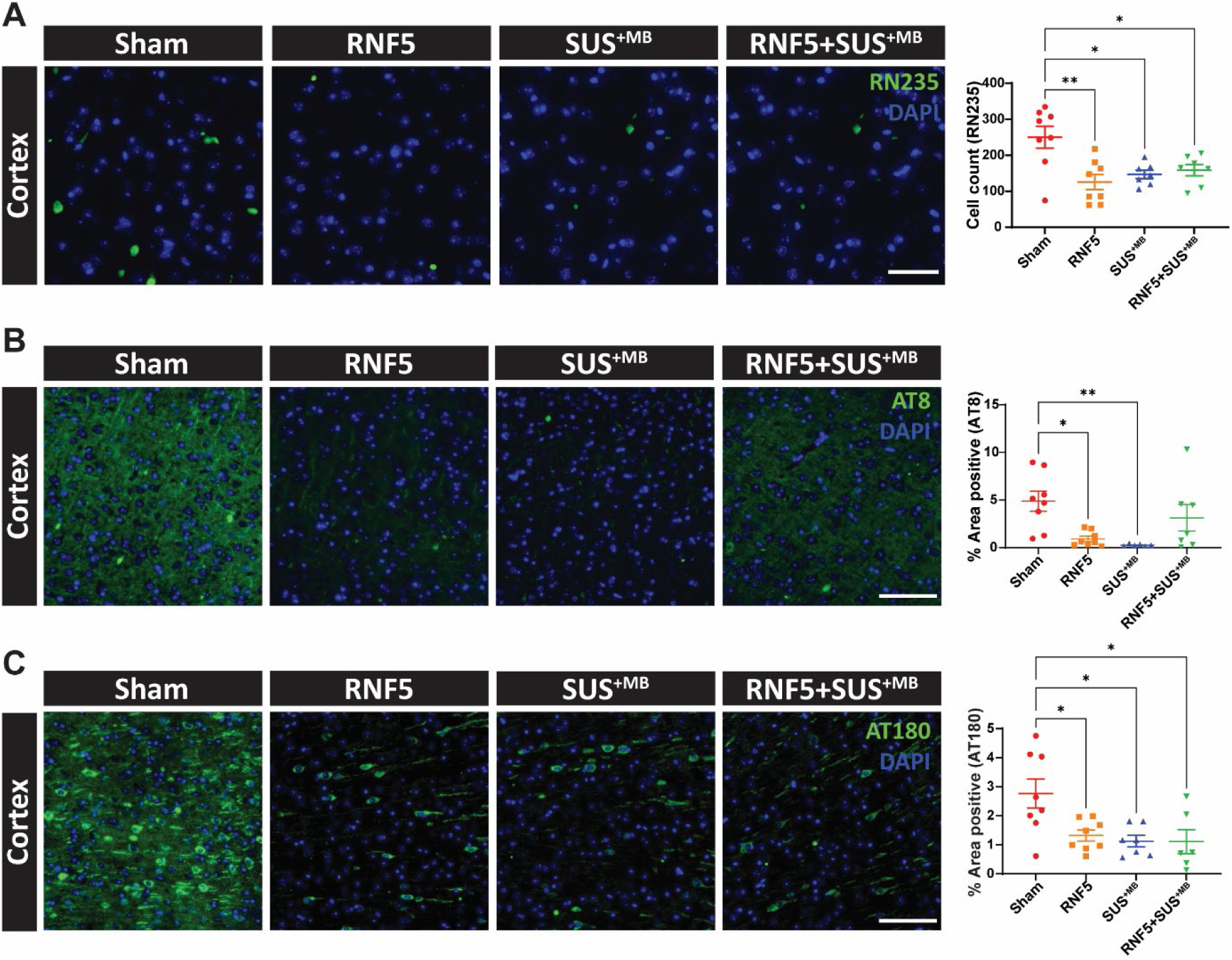
All treatments reduce phosphorylated tau in the cortex. **(A)** Representative images (20x) of RN235 (pS235) immunoreactivity observed in the cortex of K3 mice in each treatment group with quantification of RN235-positive cells. Scale bar, 50 µm. Sham, n=8; RNF5, n=8; SUS^+MB^, n=7; RNF5+SUS^+MB^, n=7. **(B)** Representative images (20x) of AT8 (pS202/pT205) immunoreactivity observed in the cortex of K3 mice in each treatment group with quantification of % area positivity of AT8-positive pick body inclusions. Scale bar, 100 µm. Sham, n=8; RNF5, n=8; SUS^+MB^, n=6; RNF5+SUS^+MB^, n=7. **(C)** Representative images (20x) of AT180 (pT231) immunoreactivity observed in the cortex of K3 mice in each treatment group with quantification of % area positivity of AT180-positive NFT-like inclusions. Scale bar, 100 µm. Sham, n=8; RNF5, n=8; SUS^+MB^, n=7; RNF5+SUS^+MB^, n=6. p<0.05, **p<0.01, ***p<0.001. All data represented as mean ± SEM; one-way ANOVA with Tukey’s multiple comparisons test.

**Figure 4.**
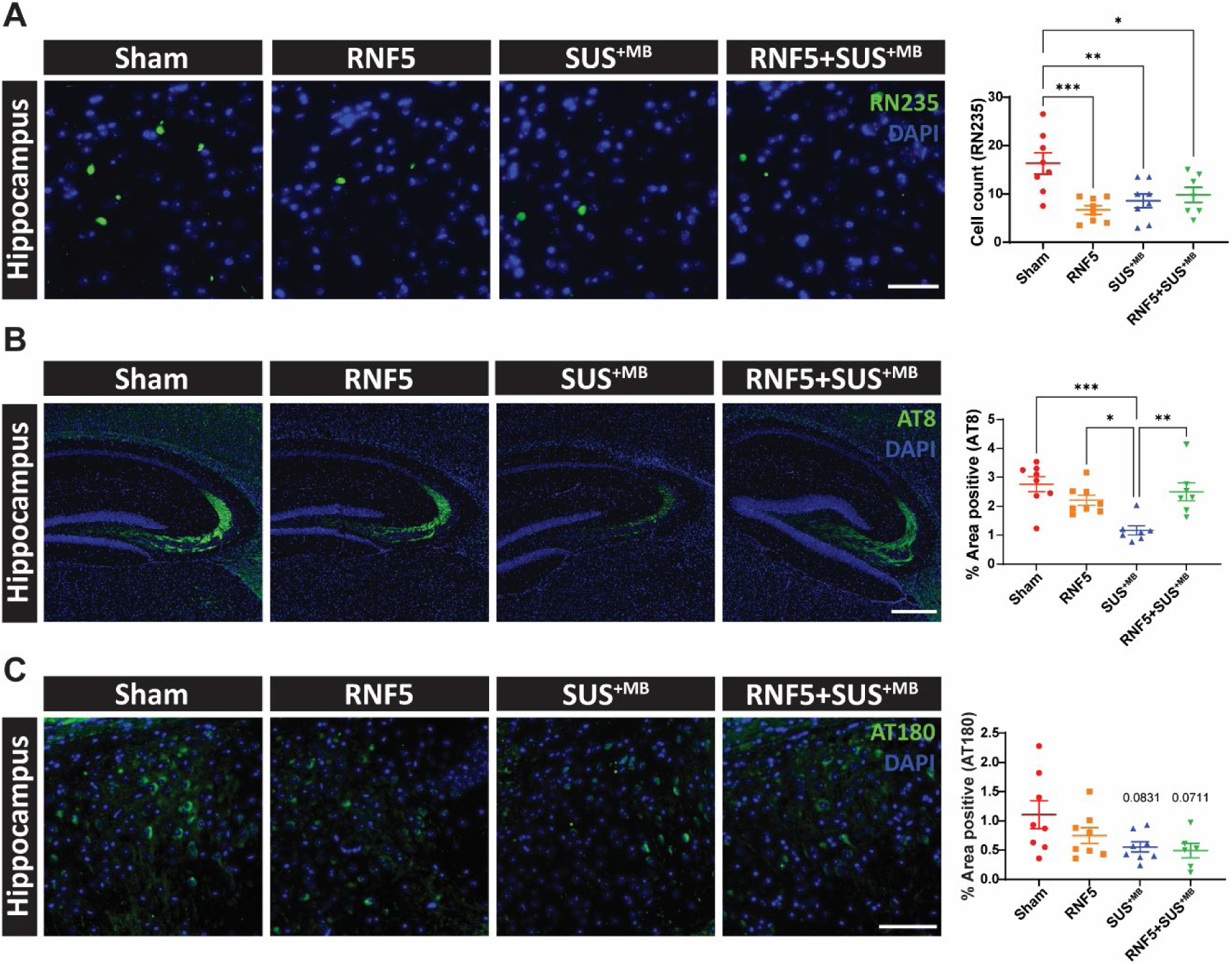
All treatments reduce phosphorylated tau in the hippocampus. **(A)** Representative images (20x) of RN235 (pS235) immunoreactivity observed in the hippocampus of K3 mice in each treatment group with quantification of RN235-positive cells. Scale bar, 50 µm. Sham, n=8; RNF5, n=8; SUS^+MB^, n=8; RNF5+SUS^+MB^, n=7. **(B)** Representative images (20x) of AT8 (pS202/pT205) immunoreactivity observed in the hippocampus of K3 mice in each treatment group with quantification of % area positivity of diffuse AT8-positive staining. Scale bar, 400 µm. Sham, n=8; RNF5, n=8; SUS^+MB^, n=7; RNF5+SUS^+MB^, n=7. **(C)** Representative images (20x) of AT180 (pT231) immunoreactivity observed in the hippocampus of K3 mice in each treatment group with quantification of % area positivity of AT180-positive NFT-like inclusions. Scale bar, 100 µm. Sham, n=8; RNF5, n=8; SUS^+MB^, n=8; RNF5+SUS^+MB^, n=6. *p<0.05, **p<0.01, ***p<0.001. All data represented as mean ± SEM; one-way ANOVA with Tukey’s multiple comparisons test.

### Comparison of total and phosphorylated tau reductions in whole brain lysates for the different treatment groups

To complement the above data, we next assessed the levels of total and phosphorylated tau in sarkosyl-soluble and insoluble fractions of whole brain lysates (excluding the cerebellum). Sarkosyl extraction is an established method to investigate soluble and insoluble (aggregated) tau in brain. In the sarkosyl-soluble fractions, which contain soluble tau, no differences were found for total mouse and human tau as detected with antibody Tau-5; also, no differences were evident in AT8 phosphorylation between any groups (Figure 5A). Detection of AT180 positive tau, however, revealed a trend towards a reduction in RNF5-treated mice compared to sham mice (Figure 5A). Most interestingly, levels of sarkosyl-soluble AT180-tau were significantly increased in the combination RNF5+SUS^+MB^ group, compared to mice treated with RNF5 only (Figure 5A). In the sarkosyl-insoluble fraction, which contains aggregated species of tau, no differences were found for total tau (Tau-5) or phospho-tau (AT8 and AT180) between any groups (Figure 5B).

**Figure 5.**
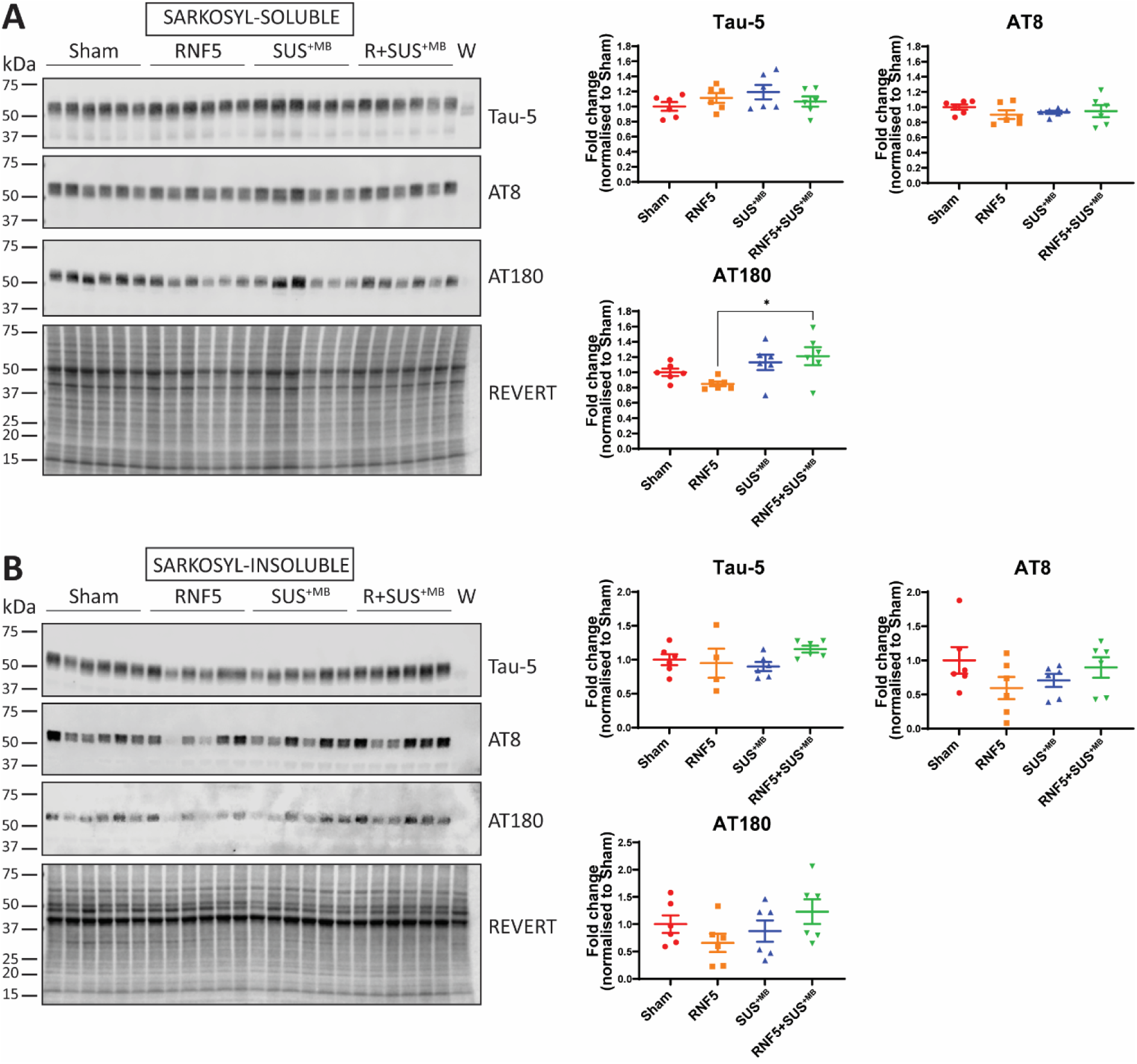
RNF5 when delivered by SUS^+MB^ does not reduce total or phosphorylated tau levels in whole brain lysate. **(A)** Representative images of sarkosyl-soluble K3 whole brain extracts probed with anti-tau antibodies as indicated and quantification. **(B)** Representative images of sarkosyl-insoluble K3 whole brain extracts probed with anti-tau antibodies as indicated and quantification (W = brain homogenate from WT mice). *p<0.05. All data represented as mean ± SEM (Sham, n=6; RNF5, n=6; SUS^+MB^, n=6; RNF5+SUS^+MB^, n=6); one-way ANOVA with Tukey’s multiple comparisons test.

### Microglial number and percentage area are elevated after the combination treatment but not after RNF5 treatment alone

Several *in vitro* and *in vivo* studies have demonstrated that antibody-tau complexes can be phagocytosed by microglia through the interaction of the antibody Fc region with Fc receptors on microglia [20, 21]. We have also previously reported that SUS^+MB^ enhances microglial activation in APP23 mice and this in turn aided plaques to be cleared through microglial phagocytosis [6]. Taken together, we speculated that the lack of enhanced efficacy in the combination group may be due to overactivation of microglia. Therefore, we sought to investigate whether microglial activation was enhanced in the combination treatment cohort compared to mice administered with the single treatments, by staining for Iba1, a cytoplasmic microglial marker (Figure 6A). We found that ‘microglial count’ and ‘microglial percentage area’, a proxy of microglial activation, were elevated in mice that had been treated with the combination compared to WT, Sham and RNF5-treated mice, but not compared to SUS^+MB^ mice (Figure 6C-E), suggesting that the SUS^+MB^ component induces microglial activation. In addition, the microglia in mice treated with RNF5 were comparable to sham treated mice despite observing significant reductions in p-tau in RNF5-treated mice compared to sham. We speculated that this elevation of Iba1-positive microglia in the groups treated with SUS^+MB^ and RNF5+SUS^+MB^ may indicate increased phagocytic activity; however, no significant differences were seen in the area of immunoreactivity for the microglial lysosomal marker CD68 between any groups (Figure 6B and F). Additionally, we observed no histological damage in any of the treated mice as shown by hematoxylin and eosin (H&E) staining (not shown). Overall, in the absence of a difference in the microglia between the SUS^+MB^ only and RNF5+SUS^+MB^ treated mice, the effects on microglia seen in the combination group are most likely mediated by the SUS^+MB^ treatment. Furthermore, these results may suggest a microglia-independent mechanism of RNF5-mediated tau pathology reduction or the possibility of an immediate short response following delivery that has resolved itself over time, which could explain the inability to detect this effect. However, since these mice were sacrificed one week following their final treatment, what we have observed in these mice so far may not be reflective of the mechanisms taking place in the hours immediately following treatment. Therefore, to gain a deeper understanding of the events immediately after delivery, mice were treated with fluorescently labelled RNF5, with or without SUS^+MB^, and sacrificed at earlier time-points.

**Figure 6.**
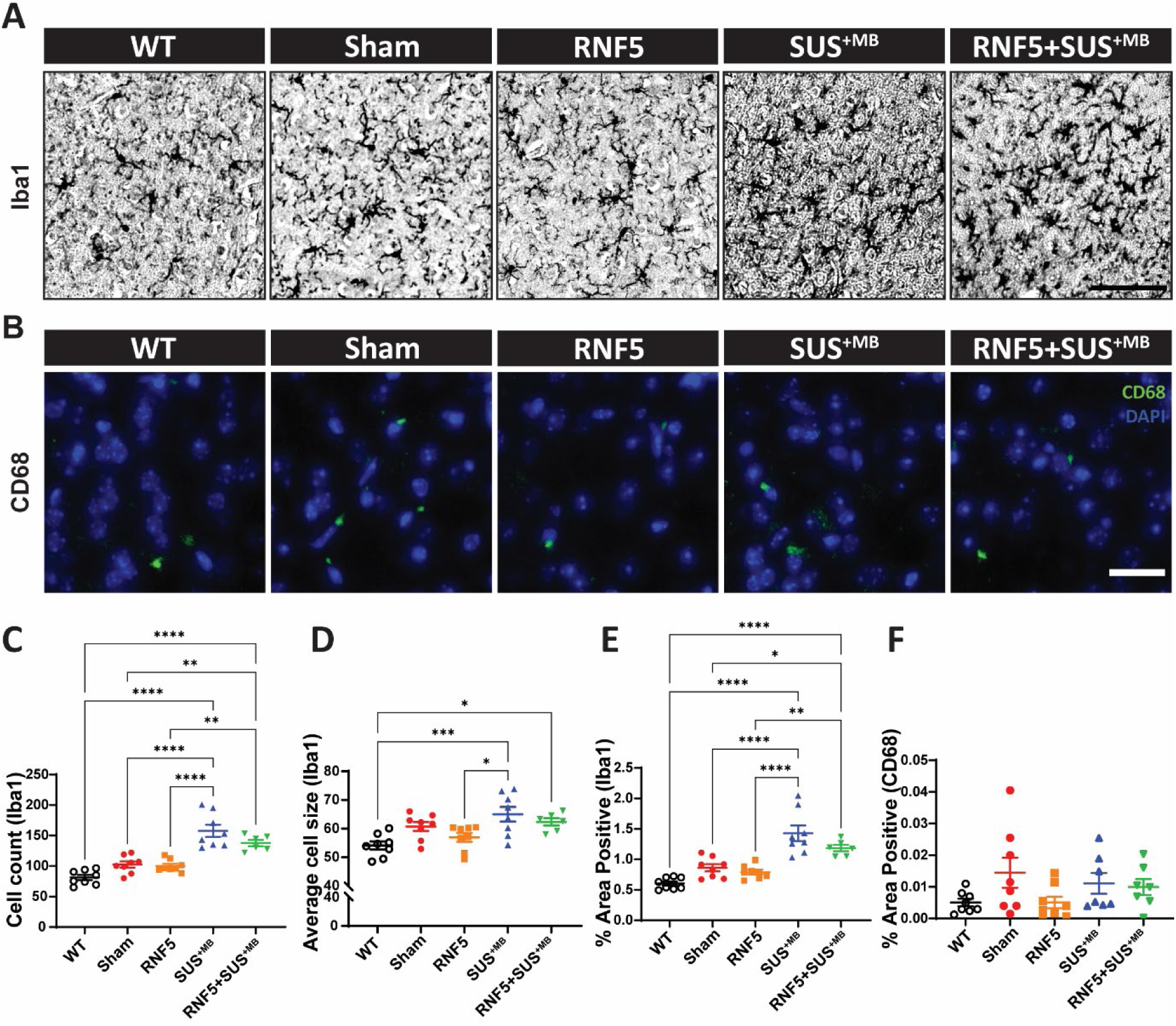
Microglia number and percentage area are elevated after combination RNF5+SUS^+MB^ treatment but not after RNF5 treatment alone. **(A)** Representative images (20x) of Iba1 pan-microglial staining in the cortex. Scale bar, 100 µm. **(B)** Representative images (20x) of CD68 microglial lysosome staining in the cortex. Scale bar, 25 µm. Quantification of (A) showing **(C)** Iba1 cell count, **(D)** Iba1 average cell size, **(E)** Iba1 % area positivity. Quantification of (B) showing **(F)** CD68 % area positivity. For Iba1 quantifications (C-E): WT, n=8; Sham, n=8; RNF5, n=8; SUS^+MB^, n=8; RNF5+SUS^+MB^, n=6; ****, p<0.0001; ***, p<0.001; **, p<0.01; *, p<0.05. For CD68 quantification (F): WT, n=8; Sham, n=8; RNF5, n=8; SUS^+MB^, n=7; RNF5+SUS^+MB^, n=7.

### SUS^+MB^ application enhances brain uptake of RNF5

We first sought to assess the localisation of fluorescently labelled RNF5 in the brain immediately after antibody delivery. To explore this, we administered fluorescently labelled RNF5 with or without SUS^+MB^ and sacrificed the mice 2 and 8 hours after treatment. At 2 hours, using an endothelial marker for detecting blood vessels (lectin) [22], RNF5 fluorescence is lower in the blood vessels when delivered without SUS^+MB^, compared to RNF5 delivered with SUS^+MB^ (Figure 7A). This is further evidenced by the fact that the brightness and contrast of the images had to be enhanced for RNF5 to be detectable in the RNF5 only group (Figure 7B). By 8 hours, RNF5 is no longer detectable when delivered without SUS^+MB^ suggesting that RNF5 may be distributed diffusely throughout the brain parenchyma at levels below detection (Figure 7B). On the other hand, when RNF5 is delivered with SUS^+MB^, while still localised to blood vessels at 8 hours, a portion of RNF5 is detectable in the brain parenchyma (Figure 7C) and in cells (Figure 7D). As RNF5 is detectable in the brain parenchyma at 8 hours following SUS^+MB^ treatment suggests increased brain penetration of RNF5 compared to mice treated with RNF5 without SUS^+MB^, as expected. However, as RNF5 is detected in the blood vessels at this time-point, it might get trapped in the vessels following treatment with SUS^+MB^, which does not occur without ultrasound. In fact, we were able to detect RNF5 in blood vessels at 24 hours post-SUS^+MB^ (not shown). Furthermore, as RNF5 was only detectable in the brain parenchyma at the 8-hour, but not the 2-hour time-point following SUS^+MB^, it suggests that there is a sequential process by which IgG traverses the BBB, rather than an immediate transfer between the brain endothelial cells in a paracellular fashion.

**Figure 7.**
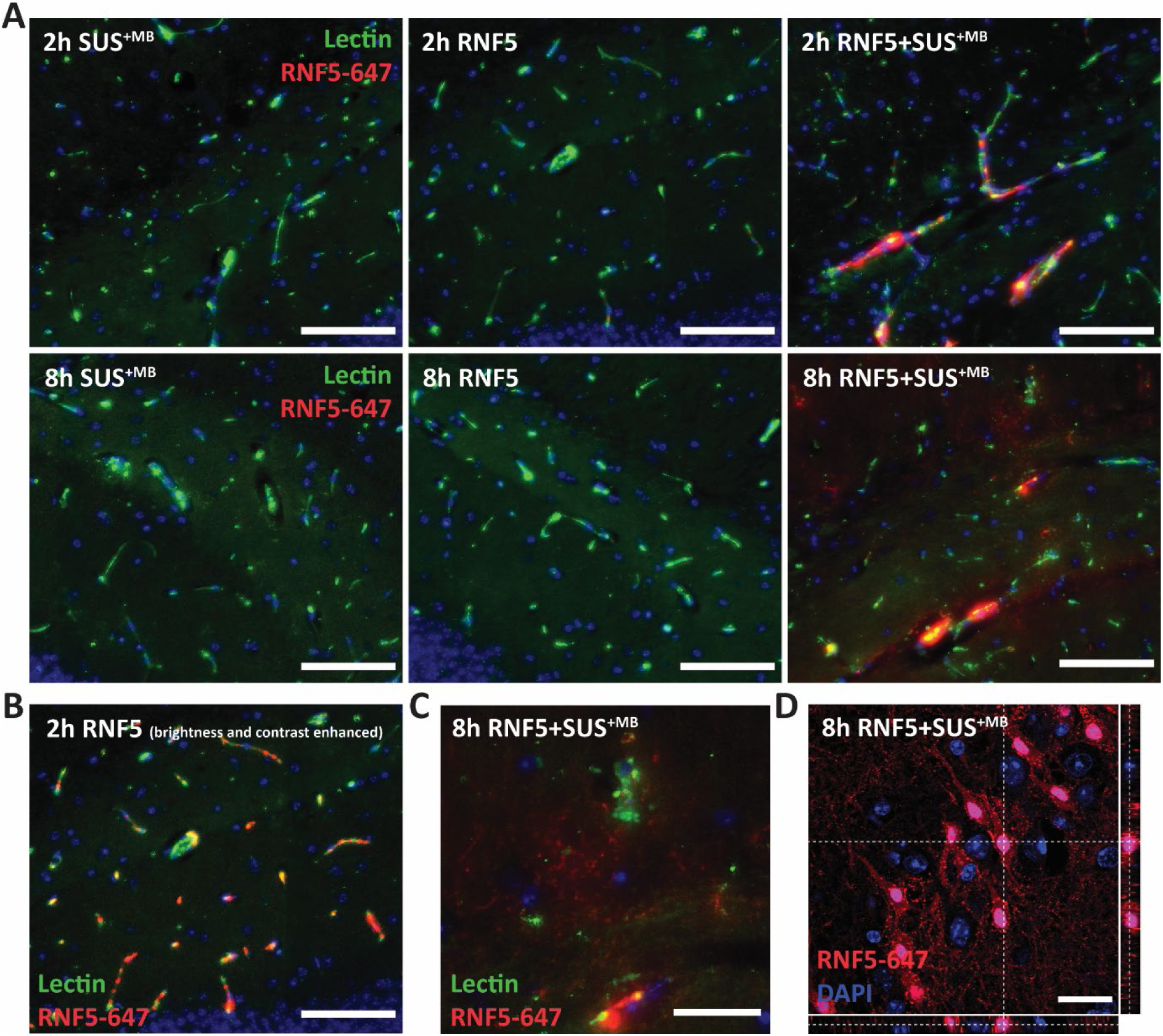
RNF5 IgG antibody localisation studies following SUS^+MB^ application. **(A-C)** Representative images (20x) of the hippocampus of mice treated with RNF5, SUS^+MB^ or RNF5+SUS^+MB^ at 2- and 8-hours post-treatment. Lectin immunoreactivity represents microvessels. **(A)** In K3 mice treated with RNF5 alone, RNF5 is below detection limits at 2 hours and 8 hours at the brightness and contrast applied in these images. In K3 mice treated with RNF5+SUS^+MB^, RNF5 is localised to microvessels at 2 hours, and by 8 hours, is still detectable in microvessels. Scale bar, 100 µm. **(B)** After brightness and contrast enhancement, RNF5 is detectable in the blood vessels of mice treated with RNF5 alone at 2-hours post-treatment. Scale bar, 100 µm. Complete figure shown in Supplementary Figure 2. **(C)** At 8-hours following combination treatment, RNF5 has also extravasated into the brain parenchyma. Scale bar, 40 µm. **(D)** Representative Z-stack images (63x) with orthogonal views showing RNF5-647 internalisation into cells in the CA3 of the hippocampus of K3 mice treated with RNF5+SUS^+MB^ at 8-hours post-treatment. Scale bar, 25 µm.

### RNF5 IgG delivery following SUS^+MB^ application induces acute neuronal damage

To determine whether RNF5 delivered with SUS^+MB^ is localised to neurons as a target cell-type with tau deposition, neuronal co-staining was conducted with MAP2. Surprisingly, we observed that the cells in the hippocampus that had internalised RNF5 at 8 hours post-treatment were MAP2-negative (Figure 8A). They were also Iba1-and GFAP-negative indicating that RNF5 was not being internalised by microglia or astrocytes in these areas either (not shown). Given that MAP2 loss is associated with neuronal degeneration as observed in animal models of stroke [23], we determined whether the lack of MAP2 in these cells may be due to poor neuronal health. Therefore, we performed H&E staining in mice sacrificed at 2 and 8 hours following RNF5+SUS^+MB^ treatment and eosinophilic neurons or ‘red’ neurons in the hippocampus were quantified. This region was analysed as a proxy for overall brain health because we had previously observed that the pyramidal cell layer in the hippocampus was particularly vulnerable to neuronal damage compared to other brain regions. Surprisingly, we observed eosinophilic staining of neurons in the pyramidal layer of the hippocampus matching up to the areas where we observed RNF5 internalisation, particularly for the 8-hour post-treatment time-point. Quantification of the percentage of eosinophilic cells out of the total cell number in the pyramidal layer of the hippocampus demonstrated a significant increase in the number of eosinophilic cells in the 8-hour group compared to the 2-hour group (Figure 8B), indicating that neuronal degeneration was occurring over time. We also observed microbleeds (small haemorrhages) at both time-points (Figure 8B).

**Figure 8.**
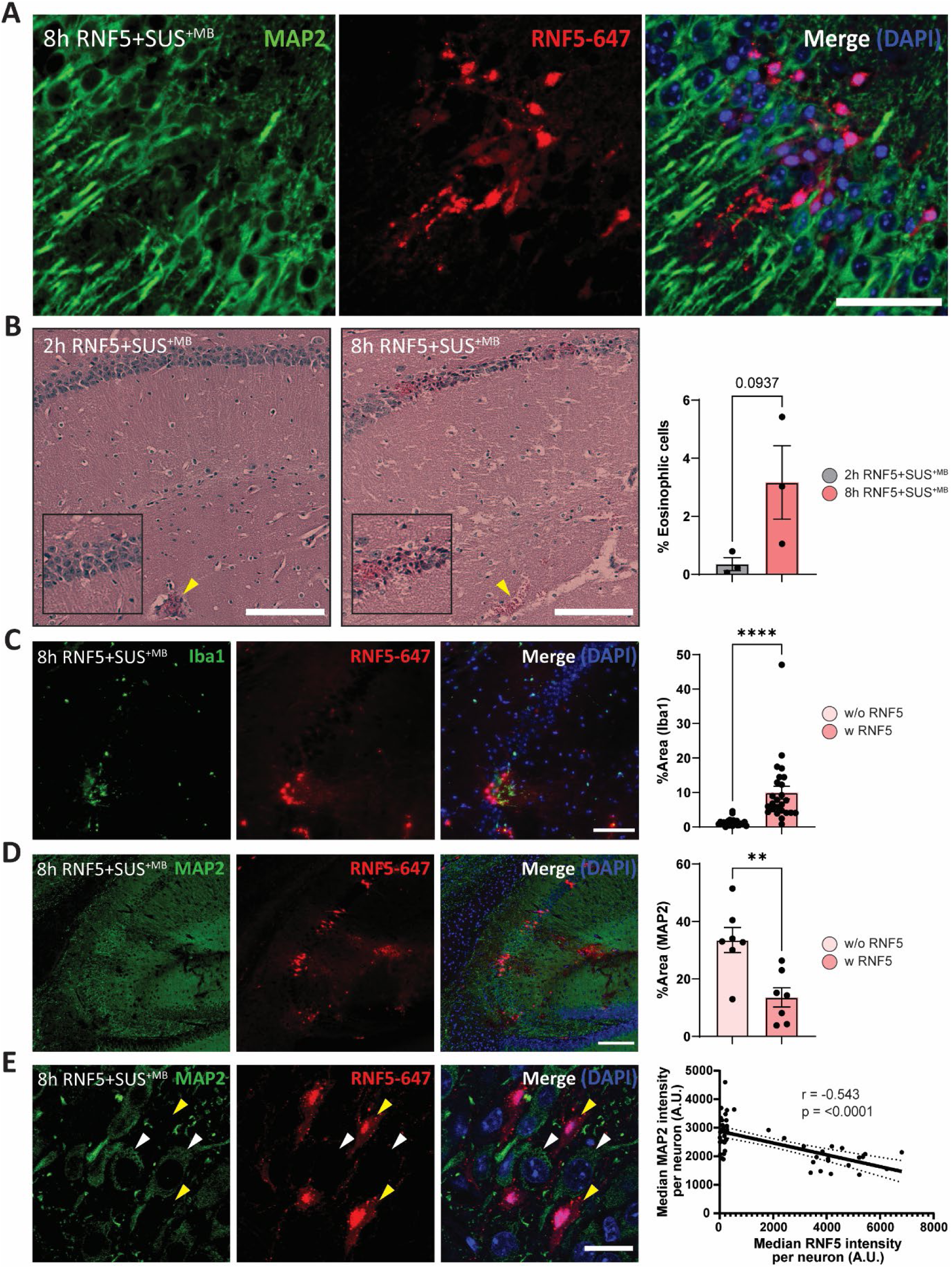
The impact of RNF5 IgG delivery following SUS^+MB^ application on brain and neuronal health. **(A)** Representative images (63x) showing maximum intensity projection of RNF5-647 internalisation into MAP2-negative neurons in the CA1 of the hippocampus of K3 mice treated with RNF5+SUS^+MB^ at 8-hours post-treatment. MAP2 immunoreactivity represents neuronal cell body and dendritic staining. Scale bar, 50 µm. **(B)** Representative images (20x) of H&E staining of K3 mice treated with RNF5+SUS^+MB^ sacrificed at 2 and 8 hours post-treatment with arrows pointing to microbleeds. Quantification of percentage of eosinophilic cells in the pyramidal layer of the hippocampus indicates a trend towards an increase in the number of eosinophilic cells in the K3 mice 8 hours post-treatment compared to 2 hours post-treatment. n=3 for each group. Scale bar, 150 µm. **(C)** Representative images (20x) of Iba1 (pan-microglia) and RNF5-647 immunoreactivity observed in the hippocampus of K3 mice treated with RNF5+SUS^+MB^ sacrificed at 8 hours post-treatment. Quantification of % area positivity of Iba1 in areas with (w) RNF5-positivity compared to areas without (w/o) RNF5-positivity indicates a significant increase in microglial infiltration into the areas where RNF5 has extravasated into the brain parenchyma/cells as well as areas around blood vessels. n=25 ROIs for each group across 3 mice. Scale bar, 100 µm. **(D)** Representative images (20x) of MAP2 and RNF5-647 immunoreactivity observed in the hippocampus of K3 mice treated with RNF5+SUS^+MB^ sacrificed at 8 hours post-treatment. Quantification of % area positivity of MAP2 in areas with (w) RNF5-positivity compared to areas without (w/o) RNF5-positivity indicates a significant reduction in neuronal health in the areas where RNF5 has extravasated into the brain parenchyma/cells. n=7 ROIs for each group across the 2 mice that had RNF5 extravasation into the parenchyma. Scale bar, 200 µm. **(E)** Representative Z-stack images (63x) showing RNF5 internalisation into MAP2-negative neurons in the CA3 of the hippocampus of K3 mice treated with RNF5+SUS^+MB^ at 8-hours post-treatment. Neurons with high MAP2 signal and low RNF5 signal are indicated by a white arrow, whereas neurons with low MAP2 signal and high RNF5 signal are indicated by a yellow arrow. Per neuron correlative analysis indicates a negative correlation between MAP2 and RNF5 (Spearman’s correlation, r = –0.543, p<0.0001, 95% confidence band shown by black dotted lines, total of 49 neurons analysed, n = 2 animals). Scale bar, 20 µm. All data represented as mean ± SEM except for correlative analysis; t-test. **p<0.01. ****p<0001.

Since the 8-hour time-point had the most detectable neuronal degeneration, we focused on this time-point for subsequent investigations. At this time-point, there was a 7-fold increase in microglial infiltration in the areas in which RNF5 had extravasated/been internalised into neurons compared to areas in which RNF5 was not present, which indicates that inflammation and possible microgliosis were occurring in areas where RNF5 was present (Figure 8C). We also observed a 2.5-fold reduction in the percentage area of MAP2 in these areas compared to areas without RNF5 (Figure 8D). To investigate this in more depth, we performed a per neuron correlative analysis and observed a negative correlation between MAP2 and RNF5 (r = –0.543, p<0.0001) indicating that neuronal degeneration is associated with RNF5 uptake (Figure 8E).

## Discussion and conclusions

Developing effective treatments for CNS disorders continues to present a major challenge due to the role of the BBB in preventing the passage of drugs, including biologics such as antibodies, across the brain. Therefore, in this study, we not only characterised the efficacy of a novel anti-tau mAb, RNF5, but also when combined with SUS^+MB^ to increase brain penetration of the antibody, anticipating an increased therapeutic effect. Our major finding was that treatment with either RNF5 or SUS^+MB^ alone, broadly speaking, reduced phosphorylated tau levels in K3 mice to a similar extent, whereas the combination treatment (RNF5+SUS^+MB^) had no additive effects, despite an increase in antibody uptake. Given that K3 mice display both tau-induced cognitive and motor impairments, we used the latter as a proxy for a potential therapeutic effect. The significant reductions in pathology however were not reflected by an overall improvement in motor impairment. We speculate that given the pronounced, progressive pathology, the achieved reductions in pathological tau were not sufficient to improve the behavioural deficits. Furthermore, given the reported degeneration of cerebellar basket cells as well as of the sciatic nerve in the K3 mice [14, 24], it is possible that SUS^+MB^ sonication did not target all of this tissue, and similarly, the tissue may not have been equally accessible to the antibody.

In the absence of SUS^+MB^, we demonstrate that intravenous injections of RNF5 alone was able to reduce tau pathology in K3 mice. This highlights that RNF5 is an effective treatment by itself. Importantly, the RNF5 signal was not observed in the brain parenchyma of mice in the absence of SUS^+MB^. This is consistent with what has been reported for other tau IgGs in clinical development [25, 26] and suggests that IgG levels may be below the detection limit but still sufficient to reduce phosphorylated tau.

It is not possible to conclusively determine the mechanism of action of RNF5 and its role in tau reduction in this study. When delivered with SUS^+MB^, RNF5 was only internalised into degenerating neurons and this is in line with other publications that have shown that antibodies are not readily internalised into neurons [25] or are only detected in dying neurons [27] which presents the possibility that RNF5 is acting in the extracellular space. Since we did not detect any increase in the phagocytic activity of the microglia in the RNF5-treated mice, we speculate that microglial phagocytosis may not be playing a significant role in tau clearance, despite being one of the well-documented mechanisms of action following passive immunotherapy with tau antibodies [20, 21, 28, 29]. Alternatively, it may be possible that RNF5 is acting in the extracellular space to prevent the internalisation of tau seeds in the K3 mice, thus preventing the formation of insoluble tau species. This is additionally supported by other reports that N-terminal tau antibodies prevented tau seeding in cell-based assays [30] as well as in passive immunisation studies in tau transgenic mice [25, 31-33].

Focused ultrasound is known to open the BBB by disrupting tight junction integrity [34] as well as increasing vesicular formation [35, 36], thus allowing the transport of large molecules across the BBB through paracellular and transcellular routes, respectively [37, 38]. While many factors dictate uptake, including size, charge and shape, a recent study has shown a role for transcytoplasmic, caveolin-mediated mechanisms in the transport of large cargoes [38]. Given that paracellular transport is likely an instant process [37] but we did not observe parenchymal localisation of RNF5 at 2 hours post-treatment, this suggests that RNF5 traversed the BBB through vesicle-mediated transcytosis. The specifics of IgG transport across the brain following ultrasound, however, has not been determined and therefore requires further investigation.

Whilst our study and others suggest that having increased IgG concentrations in the brain is beneficial, our results seem to also suggest that attempting to deliver a high localised concentration of IgG to the brain may be detrimental using the experimental conditions applied. In this study, RNF5 was delivered at a dose of 15 mg/kg (375 µg for a 25 g mouse). In comparison to other studies that have delivered antibodies retro-orbitally, administered doses were lower without ultrasound (10 mg/kg, [39]) and even lower in combination with ultrasound (40 µg [12] and 5 mg/kg [13]). This suggests the dose of RNF5 delivered into the brain may have been too high and that the benefit of SUS^+MB^ might have reached a threshold in that attempting to deliver too much antibody into the brain may then become toxic. This is especially the case because RNF5 is of a murine IgG2b isotype, and it is important to note that high concentrations of a high effector isotype antibody may ultimately be deleterious, as we have demonstrated previously [40]. By further exploring the possibility of toxicity, on top of observing acute neuronal degeneration, we also found that microglia had infiltrated the areas in the brain parenchyma into which RNF5 had extravasated. This may explain the lack of a combinatorial effect in the RNF5+SUS^+MB^ group which is supported by studies which have shown that inflammation drives tau pathology [41, 42].

Interestingly, when RNF5 was delivered with SUS^+MB^, we observed that as RNF5 internalisation into neurons increased over time, so did the extent of neuronal degeneration. IgG internalisation into neurons has been shown to occur via various mechanisms, including FcR-mediated endocytosis [28, 43-46], internalisation mediated by electrostatic interactions with the plasma membrane [47] and through lipid rafts on the surface of the plasma membrane [48]. However, neurons are not specialised to express nor do they cope well with high intracellular concentrations of IgG, as has been shown by Elmer and colleagues, where viral-mediated expression of IgG antibodies resulted in glycoprotein accumulation and subsequent neuronal degeneration [49]. While RNF5 was not expressed intracellularly in our study, it is possible that forcing large amounts of IgG into cells can be detrimental to neuronal health due to neurons being incapable of processing large quantities of glycoprotein.

Since we also observed RNF5 in blood vessels up to 24 hours following SUS^+MB^, we also speculate that neuronal degeneration and inflammation could have been induced through hypoxia-induced ischemia due to the accumulation and subsequent trapping of IgG within blood vessels. Since tau accumulation in cerebral microvasculature has been shown in human patients with AD, progressive supranuclear palsy and Lewy body dementia, as well as in the Tg2576 tau transgenic mouse model [50], it is possible that RNF5 binds to tau within brain endothelial cells (although it is not normally detected at high levels), and the resulting over-accumulation of IgG within the microvasculature is occluding the vessels, preventing the transfer of oxygen, and causing neuronal degeneration as a result of ischemia. The loss of the somatodendritic marker MAP2 seen in our study following IgG delivery with SUS^+MB^ is a phenomenon that is characteristically observed in models of stroke where a middle cerebral artery occlusion has been performed [23], and in humans who have undergone various modes of hypoxia-ischemia-related deaths [51]. In the latter study, it has been shown that the hippocampus is selectively vulnerable to oxygen deprivation compared to other brain regions, which is in line with the current study given that we saw selective neuronal damage in the hippocampus. On the other hand, this also implies that the hippocampus may be more susceptible to antibody uptake and/or the effects of SUS^+MB^. In addition, the presence of the Fc domain may play a role in IgG entrapment within microvessels. This is supported by several studies that have shown that IgG efflux from the brain to blood is mediated through neonatal Fc receptors (FcRn) expressed on both luminal (blood-side) and abluminal (brain-side) membranes of the brain endothelial cells that line cerebral blood vessels [52], which, in our study, may contribute to the presence of RNF5 in blood vessels for a long period of time following SUS^+MB^ [53-55].

In conclusion, further studies are warranted to better understand what dictates safety and efficacy of anti-tau antibody therapeutics and how this is impacted by the delivery technique. Importantly, it will also be crucial to assess whether these mechanisms are tau-dependent and the role a pre-existing pathology has. As we have previously shown that delivery of a tau antibody in a single chain format with SUS^+MB^ increases its efficacy [10], further research into the role antibody size and the presence of the antibody Fc domain have on neuronal degeneration are warranted. Further studies would also need to tease out the effect of administering different antibody doses as well as the effect of various ultrasound parameters, such as ultrasound pressure, frequency, and the biodistribution of microbubbles on neuronal degeneration.

Overall, our findings suggest that while ultrasound-mediated delivery of RNF5 IgG markedly enhances brain uptake, the localised increase in IgG concentration in the brain may be facilitating some opposing mechanisms that counteract the combinatorial therapeutic effects one would expect following combination treatment. However, given the plethora of variables involved in this study (which includes antibody concentration, antibody isotype, ultrasound parameters, microbubble distribution, and differences in blood flow and tissue oxygenation), it is difficult to pinpoint just one cause for the lack of an additive effect. While more research is required to elucidate the feasibility of this technique, this work provides an important initial understanding for the optimisation of this delivery modality.

## Abbreviations

AD: Alzheimer’s disease
IgG: immunoglobulin
SUS: scanning ultrasound
BBB: blood-brain barrier
Iba1: Ionized calcium-binding adaptor protein-1

## Credit author statement

R.B, R.M.N, and J.G designed the research protocol. R.B performed the experiments and analysed the data. E.C. provided technical assistance with experiments. R.B and R.M.N co-wrote the manuscript. All authors were involved in the discussion and interpretation of the data and in final editing.

## Acknowledgements

The authors would like to thank Dr Martina Jones from the Queensland node of the National Biologics Facility for assistance with performing the SPR experiment. We would also like to thank Wee Kiat Koh, Dr Gerhard Leinenga, Linda Cumner and Tishila Palliyaguru for their technical support and the Queensland Brain Institute animal house staff for husbandry. Imaging/Analysis was performed at the Queensland Brain Institute’s Advanced Microscopy Facility, generously supported by the Australian Government through the ARC LIEF LE100100074.

## Ethics approval

All animal experiments were conducted under the guidelines of the Australian Code of Practice for the Care and Use of Animals for Scientific Purposes and approved by the University of Queensland Animal Ethics Committee (QBI/554/17/NHMRC).

